# The two-component response regulator OrrA confers dehydration tolerance by regulating *anaKa* expression in the cyanobacterium *Anabaena* sp. strain PCC 7120

**DOI:** 10.1101/2021.08.03.454875

**Authors:** Satoshi Kimura, Miho Sato, Xingyan Fan, Masayuki Ohmori, Shigeki Ehira

## Abstract

The aquatic cyanobacterium *Anabaena* sp. strain PCC 7120 exhibits dehydration tolerance. The regulation of gene expression in response to dehydration is crucial for the acquisition of dehydration tolerance, but the molecular mechanisms underlying dehydration responses remain unknown. In this study, the functions of the response regulator OrrA in the regulation of salt and dehydration responses were investigated. Disruption of *orrA* abolished or diminished the induction of hundreds of genes in response to salt stress and dehydration. Thus, OrrA is a principal regulator of both stress responses. In particular, OrrA plays a crucial role in dehydration tolerance because an *orrA* disruptant completely lost the ability to regrow after dehydration. Moreover, in the OrrA regulon, *anaKa* encoding a protein of unknown function was revealed to be indispensable for dehydration tolerance. OrrA and AnaK are conserved among the terrestrial cyanobacteria, suggesting their conserved functions in dehydration tolerance in cyanobacteria.

## INTRODUCTION

Cyanobacteria comprise a diverse group of bacteria characterized by oxygen-evolving photosynthesis. Their photosynthetic ability enables them to inhabit almost all illuminated environments. The habitat of cyanobacteria is not limited to aquatic ecosystems, but it extends to terrestrial ecosystems including extremely arid environments such as deserts (Pointing and Belnap, 2012). Some cyanobacterial species belonging to the genera *Nostoc* and *Chroococcidiopsis* are present in dry areas, and they exhibit high tolerance to desiccation (Singh, 2018). For example, *Chroococcidiopsis* strains isolated from deserts could form colonies after 4 years of storage in a dry state, and the terrestrial strain *Nostoc commune* was revived via inoculation into medium after storage in a desiccated state for approximately 90 years (Lipman, 1941; Fagliarone *et al*., 2017).

Loss of water from cells induces various stresses and damages various cellular components including proteins, membrane lipids, and DNA (Potts, 2001; Singh, 2018). Desiccation-tolerant cyanobacteria adopt multiple strategies to survive harsh environments during desiccation. *N. commune* cells are embedded in extracellular polysaccharide (EPS), which protects cells from desiccation (Tamaru *et al*., 2005). In the aquatic cyanobacterium *Anabaena* sp. strain PCC 7120 (hereafter *Anabaena*), EPS excretion was increased by overexpression of *sigJ*, which encodes the sigma factor of RNA polymerase, and the mutant displays higher dehydration tolerance than the wild-type strain (WT) (Yoshimura *et al*., 2007). The accumulation of compatible solutes such as trehalose and sucrose, which stabilize the proteins and cellular membranes of dehydrated cells (Tapia and Koshland, 2014), represents another strategy of desiccation tolerance. *N. commune* and a *Chroococcidiopsis* strain accumulate trehalose and sucrose in response to dehydration, salt and osmotic stress (Hershkovitz *et al*., 1991; Sakamoto *et al*., 2009). *Anabaena* also utilizes both sugars as compatible solutes, and disruption of trehalose synthesis genes decreases dehydration tolerance (Higo *et al*., 2006). Moreover, *Escherichia coli* expressing the sucrose synthesis gene *spsA* from the cyanobacterium *Synechocystis* sp. strain PCC 6803 (*Synechocystis* PCC 6803) accumulates sucrose within cells and exhibits a drastic increase in survival rate after dehydration (Billi *et al*., 2002).

Oxidative damage caused by reactive oxygen species (ROS) is one of the most deleterious effects of dehydration (França *et al*., 2007). ROS generation is increased by dehydration, resulting in lipid peroxidation, denaturation of proteins through oxidative modifications, and DNA double-strand breakage. Antioxidant defense systems that suppress ROS generation and scavenge generated ROS protect cellular components from oxidation (Singh, 2018). Because most antioxidant defense systems are proteinous, avoidance of protein oxidation is a key factor for desiccation tolerance (Daly, 2009). Desiccation-resistant bacteria display lower oxidative modification of proteins than desiccation-sensitive bacteria during dehydration (Fredrickson *et al*., 2008). *Chroococcidiopsis* sp. strain CCMEE 029 is highly tolerant to desiccation, and it does not undergo protein oxidation after 1 year of incubation under dry conditions. Contrarily, *Synechocystis* PCC 6803 exhibits substantial protein oxidation, and it is incapable of surviving under the same conditions (Fagliarone *et al*., 2017).

*Anabaena* is a model organism of bacterial cellular differentiation (Flores *et al*., 2019). It has been also used to study the molecular mechanisms of responses to stresses including dehydration (Ohmori *et al*., 2001; Wang *et al*., 2002). Close relatives of *Anabaena* such as the terrestrial strain *Nostoc* sp. HK-01 exhibit high desiccation tolerance, and *Anabaena* itself displays moderate dehydration tolerance (Katoh *et al*., 2003; Singh *et al*., 2013). Because genetic engineering cannot be performed in terrestrial *Nostoc* strains, *Anabaena* serves as a model organism for investigating the molecular mechanisms of desiccation tolerance in the terrestrial strains (Yoshimura *et al*., 2007; Xu *et al*., 2020). In *Anabaena, orrA*, which encodes a response regulator of the bacterial two-component regulatory system, was identified as a regulator of *lti2* in response to salt and osmotic stresses (Schwartz *et al*., 1998). We previously indicated that OrrA regulates the expression of *spsA, susA*, and *susB*, which participate sucrose synthesis, in response to salt stress (Ehira *et al*., 2014). Sucrose accumulation under salt stress conditions is lowered by the *orrA* disruption, and intracellular sucrose levels are increased by *orrA* overexpression (Ehira *et al*., 2014). In this study, we analyzed the global effects of *orrA* disruption on gene expression, observing that OrrA is a master regulator of dehydration and salt stress responses in *Anabaena*. Moreover, among hundreds of genes regulated by OrrA, *anaKa*, which encodes a protein of unknown function, proved essential for dehydration tolerance in *Anabaena*.

## RESULTS

### OrrA is a major regulator of salt stress response

Sucrose synthesis under salt stress conditions is regulated by OrrA in *Anabaena* (Ehira *et al*., 2014). The functions of OrrA in global salt stress response regulation were investigated by transcriptome analysis. The gene expression profiles of WT and the *orrA* mutant strain DRorrAS were compared between cells subjected to salt stress (50 mM NaCl) for 3 h and cells before stress treatment. The transcript levels of 298 genes were increased at least twofold by salt stress in WT, while those of another 1,358 genes were decreased (Table S1). Three genes involved in sucrose metabolism, namely *spsA, susA* and *susB*, were upregulated by salt stress as previously reported (Ehira *et al*., 2014). The upregulated genes included *mth* encoding an enzyme for trehalose synthesis (Higo *et al*., 2006) and *anaK* genes (all4050, all4051, and all5315), which are predominantly expressed in cyanobacterial resting cells called akinetes in *Anabaena variabilis* (Zhou and Wolk, 2002). Genes encoding proteins that prevent ROS generation, such as *hli/scp* genes (asl0514, asl0873, asr3042, and asr3043) and *dps* genes (all0458 and all1173), and genes that detoxify ROS, such as *katA* and *katB*, were also upregulated (Howe *et al*., 2018; Tibiletti *et al*., 2018). The downregulated genes included genes involved in photosynthesis (*psa* and *psb* genes) and carbon fixation (*rbcLXS, prk, ccmNMLK, ecaA*, and *cmpABCD*), and genes encoding components of the phycobilisome (*apc, cpc*, and *pec* genes). Moreover, the transcript levels of genes involved in transcription and translation (*rpl* and *rps* genes, *rpoA, rpoC1*, and *rpoC2*), and ATP synthesis (*atp* genes) were decreased. The suppression of cell growth under salt stress conditions leads to downregulation of these genes (Ehira *et al*., 2014).

In DRorrAS, the numbers of genes upregulated and downregulated by salt stress were decreased to 71 and 51, respectively (Table S1). Meanwhile, 37 genes were upregulated in both WT and DRorrAS (Fig. 1), but the degree of upregulation was higher in WT than in DRorrAS. Temporal changes in the expression of salt-induced genes were examined by quantitative reverse transcription (qRT)-PCR (Fig. 2). In WT, *mth* expression was induced within 30 min after NaCl addition, with the *mth* transcript level increasing approximately 800-fold after 60 min, but its induction was completely abolished in DRorrAS (Fig. 2A). The induction patterns of two *dps* genes (all0458 and all1173) differed from each other, but both genes lost salt-responsive induction by *orrA* disruption (Fig. 2B). There are four paralogs of *anaK* (*anaKa, anaKb, anaKc*, and *anaKd*) in *Anabaena* (Zhou and Wolk, 2002). All *anaK* genes were induced by salt stress, which depended on *orrA* (Fig. 2C). These results indicate that OrrA is a major regulator of salt stress response in *Anabaena*.

**Fig 1.**
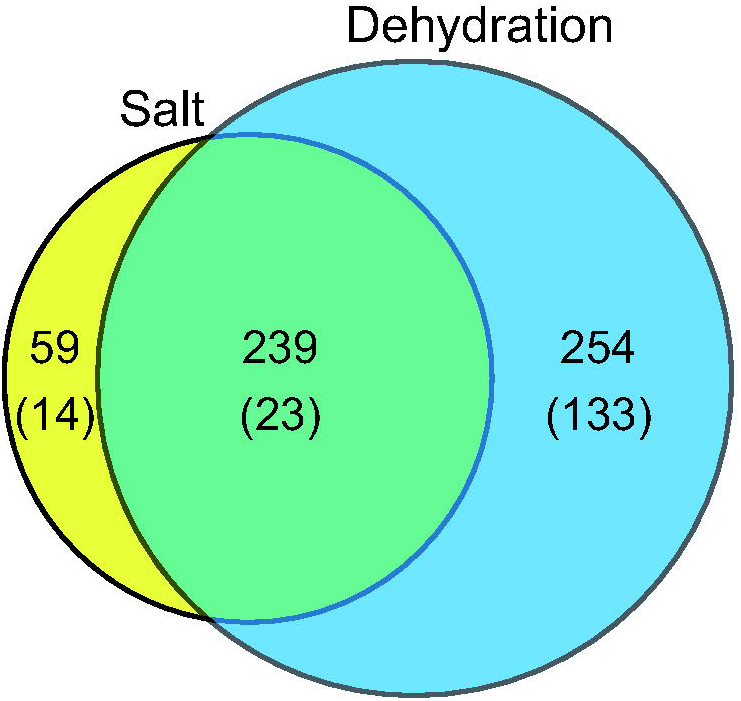
Venn diagrams of genes upregulated by salt stress and dehydration in wild-type *Anabaena* sp. strain PCC 7120 (WT) and the *orrA* disruptant DRorrAS. The numbers in parenthesis indicate genes upregulated in both WT and DRorrAS.

**Fig 2.**
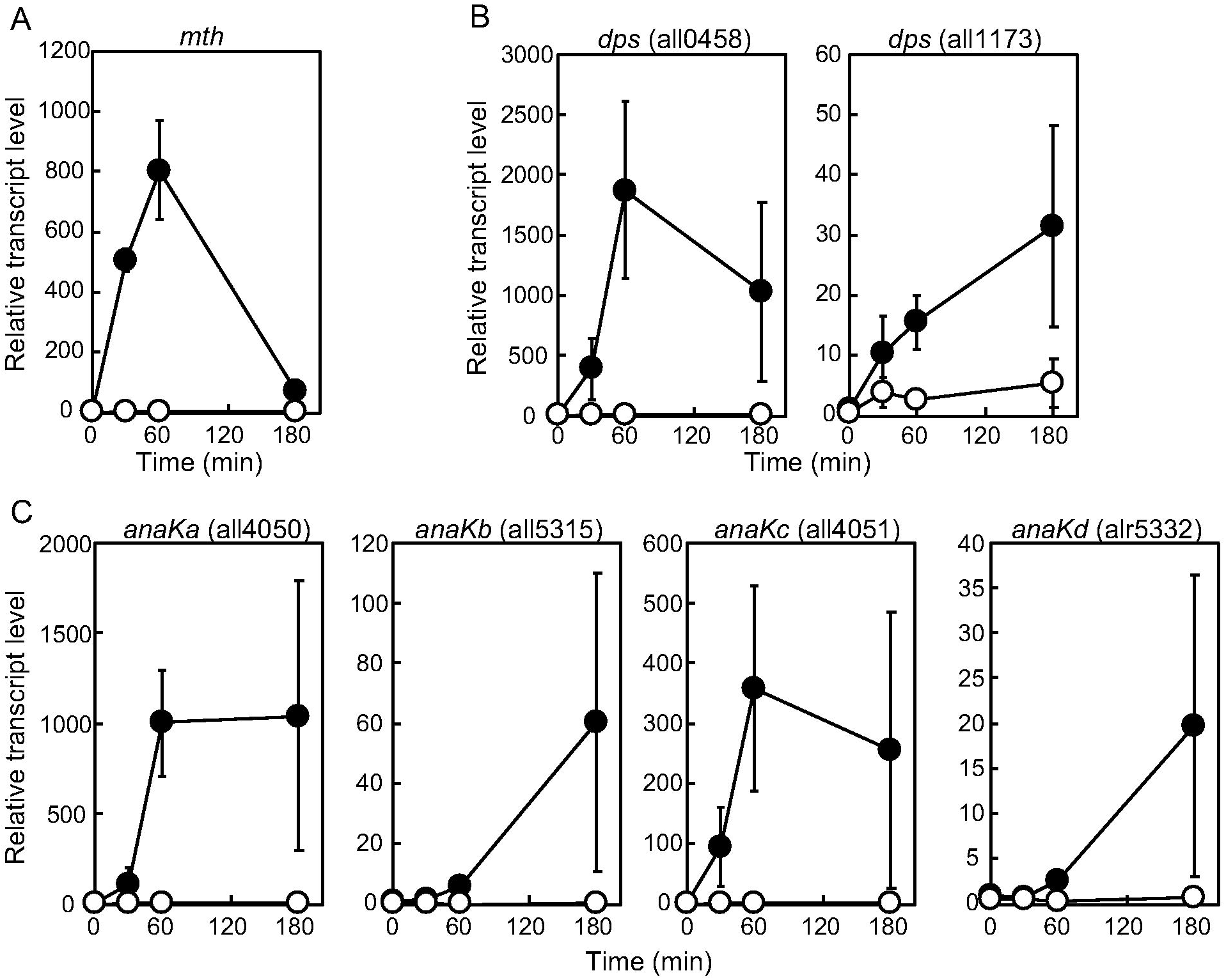
Temporal changes in the transcript levels of genes upregulated by salt stress. The transcript levels of *mth* (A), *dps* (B), and *anaK* (C) after the addition of 50 mM NaCl were determined by qRT-PCR in WT (black circles) and the *orrA* disruptant (white circles). The transcript levels were determined in duplicated measurements using three independently grown cultures. The transcript level at 0 min in WT was taken as 1. Data are presented as the mean ± SD.

### orrA *is necessary for dehydration tolerance*

OrrA regulates the expression of most salt-responsive genes, whereas its disruption does not affect growth under salt stress conditions (Ehira *et al*., 2014). Expression of *orrA* is induced by not only salt stress, but also dehydration (Higo *et al*., 2006). Moreover, many genes upregulated by salt stress, such as *mth, hli/scp* genes, and *dps* genes, are also upregulated by dehydration (Higo *et al*., 2006). To investigate the effect of the *orrA* disruption on gene expression in response to dehydration, gene expression profiles were compared between WT and DRorrAS after 3 h of dehydration. In WT, 493 genes were upregulated by dehydration, while 1,307 genes were downregulated (Table S1). Among the upregulated genes in WT, 156 genes were also upregulated in DRorrAS, but the extent of upregulation was significantly lower in DRorrAS than in WT (Fig. 1 and Table S1). Salt stress and dehydration increased the expression of the same 239 genes, only 23 of which were upregulated by both stresses in DRorrS (Fig. 1). *spsA, susA*, and *susB*, which are involved in sucrose metabolism, were also induced by dehydration under the control of OrrA (Table S1). The intracellular sucrose level in WT was increased more than 80-fold after 9 h of dehydration, whereas the level was twofold lower in DRorrAS than in WT (Table 1). Thus, OrrA plays a principal role in regulating gene expression during dehydration in *Anabaena*.

**Table 1.** Sucrose contents after dehydration

| Strain | Sucrose ($\mu\text{mol/ mg chl}a$) | |
| --- | --- | --- |
|  | 0 h | 9 h |
| WT | $0.06 \pm 0.06$ | $5.00 \pm 0.39$ |
| DRorrAS | $0.04 \pm 0.04$ | $2.26 \pm 0.29$ |

WT, wild-type *Anabaena* sp. strain PCC 7120; DRorrAS, *orrA* mutant *Anabaena* sp. strain PCC
7120; chl $a$ , chlorophyll $a$

To investigate the physiological role of OrrA in dehydration tolerance, growth ability after dehydration was analyzed. *Anabaena* filaments grown in the liquid medium were collected by filtration, and then filaments on the filters were dried in an incubator. After the indicated times, filaments were resuspended in liquid medium and incubated for 3 days. The weight of the filters decreased during the first 8 h of incubation before reaching a plateau, indicating that all water had evaporated (Fig. 3A). In WT, filaments were capable of restarting growth even after 24 h of incubation on the filters (Fig. 3B). DRorrAS filaments were able to regrow after 6 h of incubation on the filters, but their growth was severely impaired after 7 and 8 h of incubation (Fig. 3B). Moreover, DRorrAS completely lost its ability to grow after more than 10 h of incubation on the filters. Thus, *orrA* is necessary for survival under dehydration conditions.

**Fig 3.**
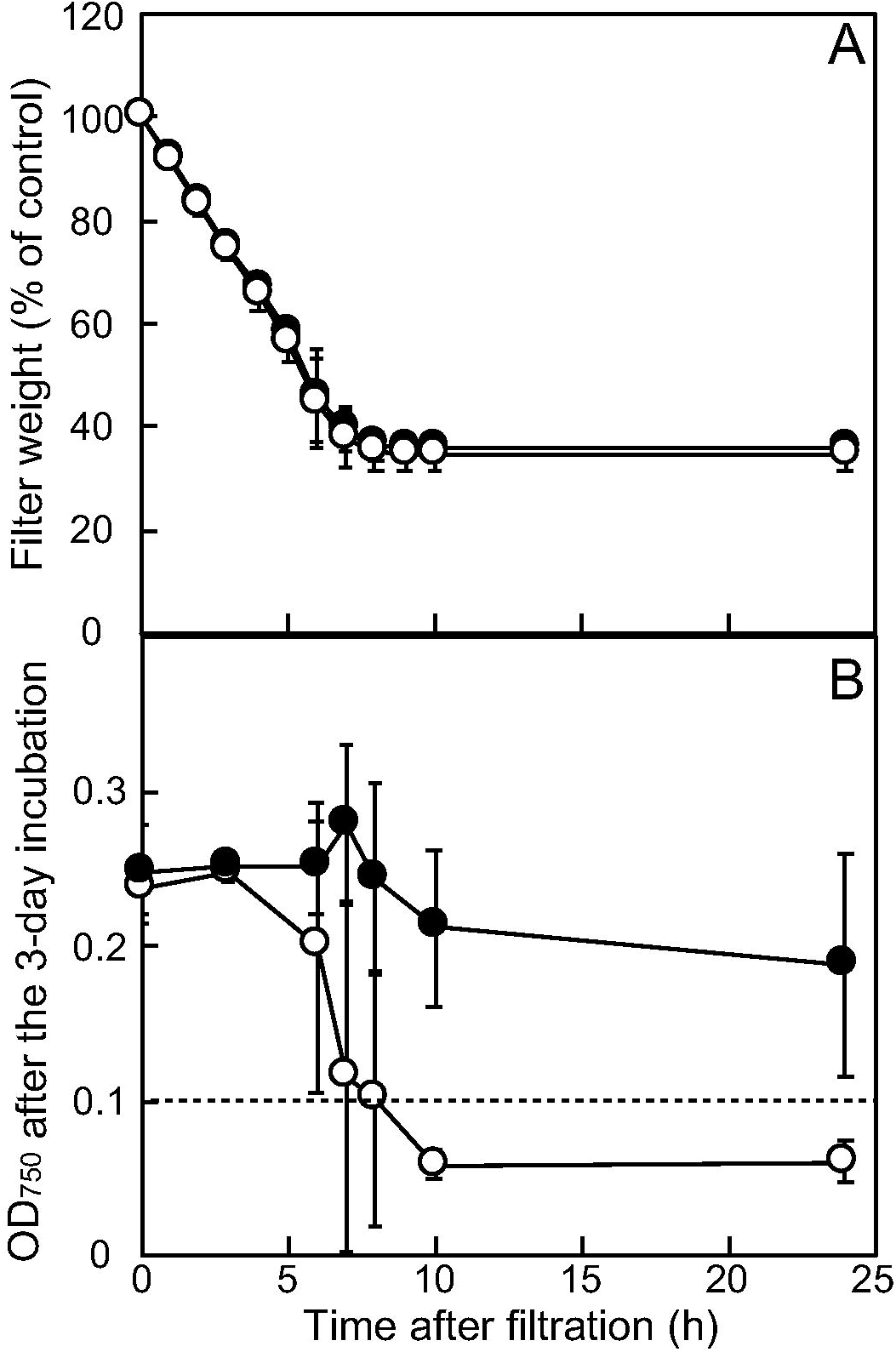
Dehydration tolerance of the *orrA* disruptant. A. Changes in weight of filters during 24h of incubation. Filaments of WT (black circles) and the *orrA* disruptant (white circles) were collected on filters and the filters dried for 24 h. The weight of each filter was measured during the 24-h incubation and presented as a ratio to the control (0 h). B. Growth ability of the *orrA* disruptant after dehydration. Filaments on the filters were resuspended in liquid medium at the indicated time, the optical density at 750 nm (OD_750_) of the suspension was adjusted to 0.1, and then filaments were incubated for 3 days in liquid medium. OD_750_ after the 3-day incubation was indicated for WT (black circles) and the *orrA* disruptant (white circles). The experiments were repeated using three independently grown cultures. Data are presented as the mean ± SD.

### orrA *is also involved in cold acclimation*

OrrA was originally identified as a regulator of the salt and osmolyte responses of *lti2* (Schwartz *et al*., 1998). In this study, we confirmed the OrrA-regulated *lti2* expression in response to salt and dehydration (Table S1). *lti2* is also known to be upregulated under low temperatures (Ehira *et al*., 2005). In addition, low temperature-responsive genes, such as alr0169, alr0803, alr0804, and alr1819, were under the control of OrrA (Table S1). Hence, we analyzed the effects of *orrA* disruption on cold acclimation. *orrA* expression increased within 30 min after the temperature was decreased from 30°C to 22°C (Fig. 4A). Growth of DRorrAS at 30°C is comparable to that of WT (Ehira *et al*., 2014), but its growth at 22°C was slower than that of WT (Fig. 4B). Thus, OrrA also influences cold accilimation.

**Fig 4.**
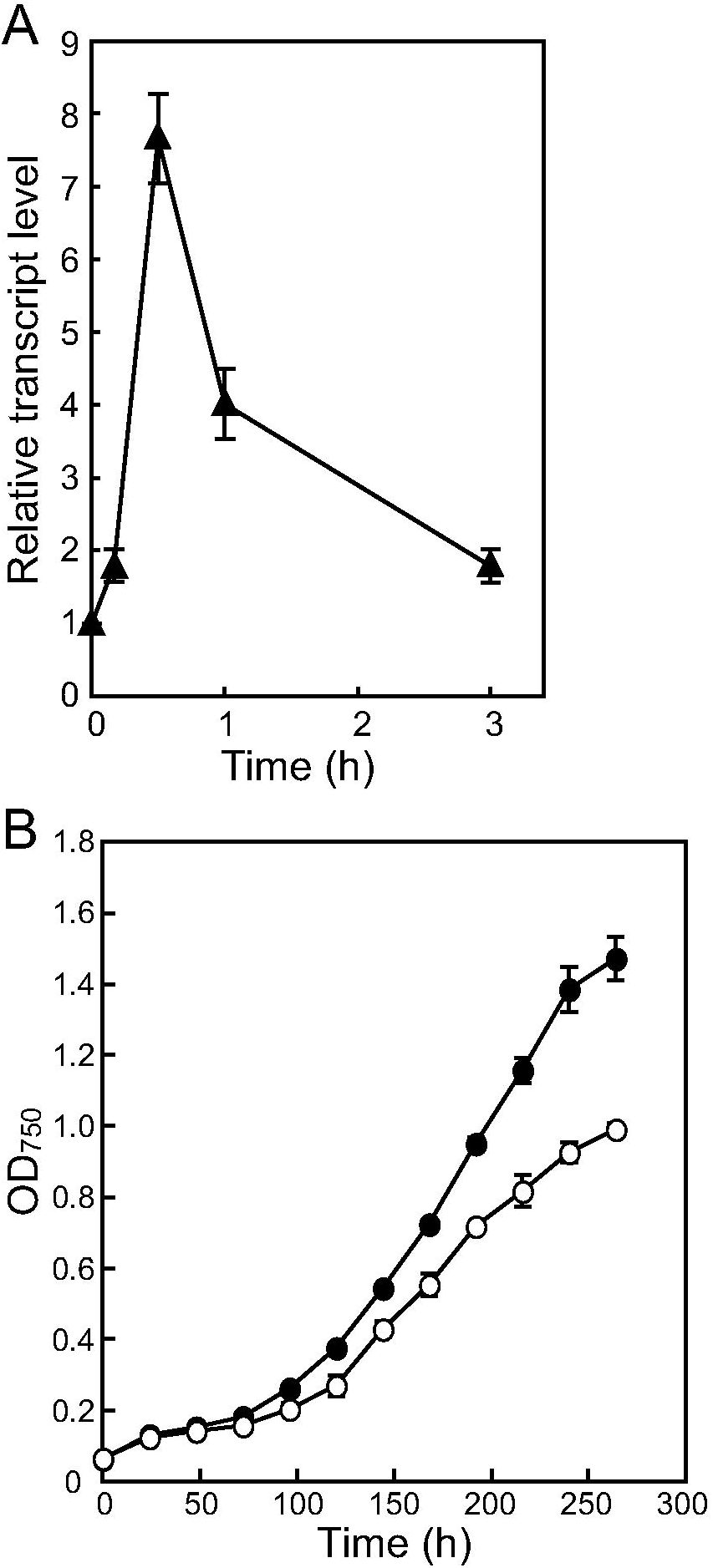
Involvement of OrrA in cold acclimation. A. Changes in the *orrA* transcript levels after a shift to low temperature (22°C). The transcript level at 0 h was taken as 1. B. Growth of WT (black circles) and the *orrA* disruptant (white circles) at 22°C. Growth was monitored by measuring OD_750_. The experiments were repeated with three independently grown cultures. Data are presented as the mean ± SD.

### anaKa *is a determinant of dehydration tolerance*

To further reveal the mechanisms of dehydration tolerance, the involvement of genes regulated by OrrA in dehydration tolerance was investigated. We focused on *anaK* because this gene is highly expressed in akinetes (Zhou and Wolk, 2002), which are cells that are extremely resistant to stresses such as cold and desiccation (Adams and Duggan, 1999). Four *anaK* paralogs, namely *anaKa, anaKb, anaKc*, and *anaKd*, were induced by dehydration ,and this induction depended on OrrA (Table S1). Each *anaK* paralog was inactivated, and the growth of the disruptants after dehydration was determined (Fig. 5). The *anaKa* disruptant DRanaKaK could not grow after incubation on the filters for 24 h, whereas the other *anaK* disruptants retained similar growth ability as WT (Fig. 5A). Time-course experiments illustrated that DRanaKaK lost growth ability at 8 h (Fig. 5B), when water had completely evaporated from filters (Fig. 3A). This indicated that *anaKa* disruption has a fatal effect on survival under dehydration.

**Fig 5.**
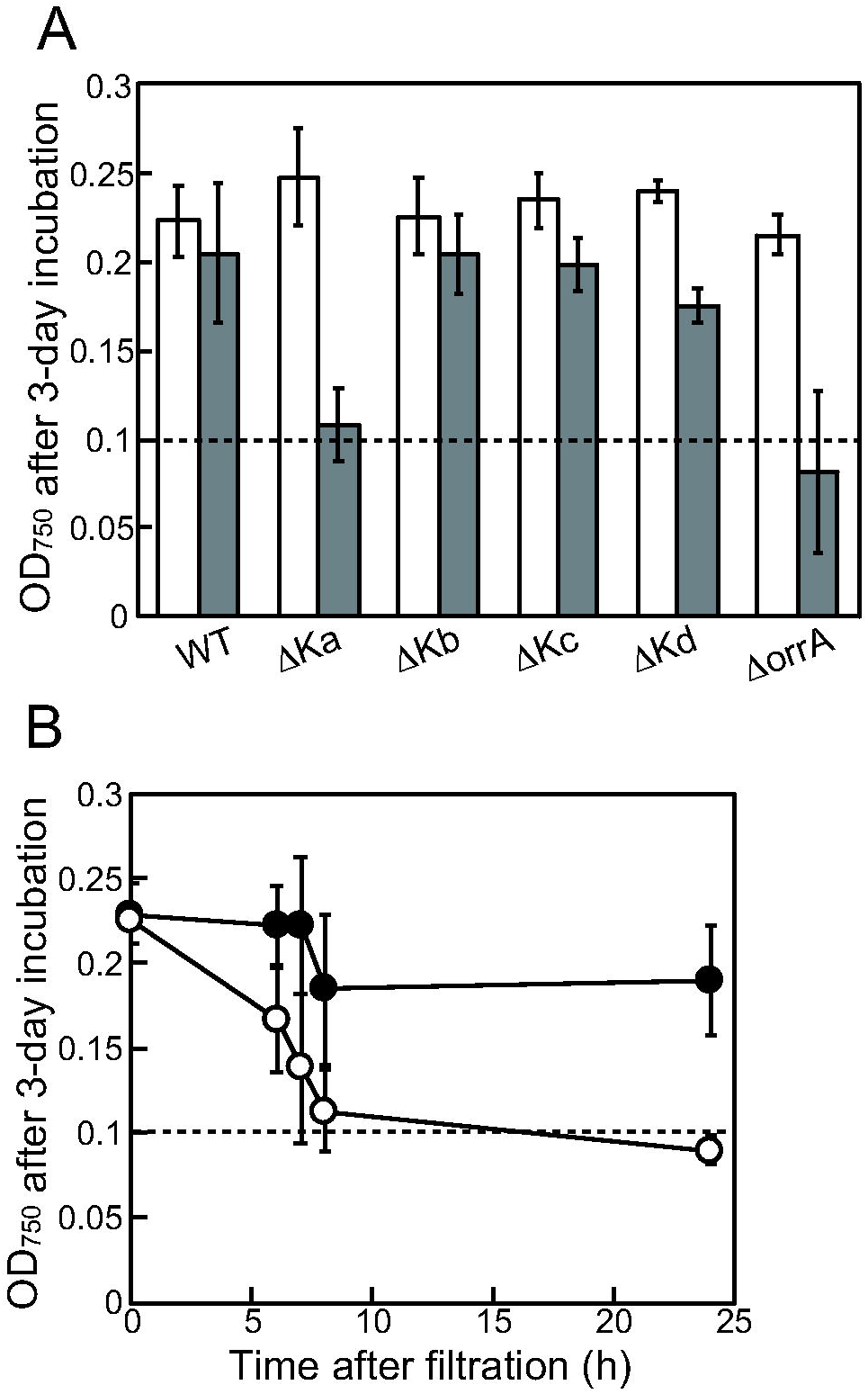
Dehydration tolerance of disruptants of the *anaK* genes. Filaments of disruptants of *anaKa* (ΔKa), *anaKb* (ΔKb), *anaKc* (ΔKc), *anaKd* (ΔKd), and *orrA* (ΔorrA) were subjected to dehydration for 0 (white bars) or 24 h (gray bars), and then their growth ability was evaluated as described in Fig 3 (A). Temporal changes of the growth ability of WT (black circles) and the *anaKa* disruptant (white circles) during 24 h of incubation were determined (B). The experiments were repeated using three independently grown cultures. Data are presented as the mean ± SD.

Disruption of *anaKa* alone resulted in the loss of dehydration tolerance. To examine the involvement of other genes regulated by OrrA in dehydration tolerance, damage of DRorrAS and DRanaKaK cell after dehydration was evaluated by quantifying phycocyanin excretion from cells (Fig. 6). Phycocyanin is a water-soluble pigment protein and is released extracellularly upon cell lysis (Arii *et al*., 2015). In WT, phycocyanin excretion was not enhanced by dehydration. In DRorrAS and DRanaKaK, excreted phycocyanin levels were comparable to WT levels before dehydration, but phycocyanin excretion was increased by dehydration. Phycocyanin excretion was more than twofold higher in DRorrAS than DRanaKaK, indicating greater cell membrane damage in DRorrAS. Thus, the *orrA* disruption is more detrimental than the *anaKa* disruption.

**Fig 6.**
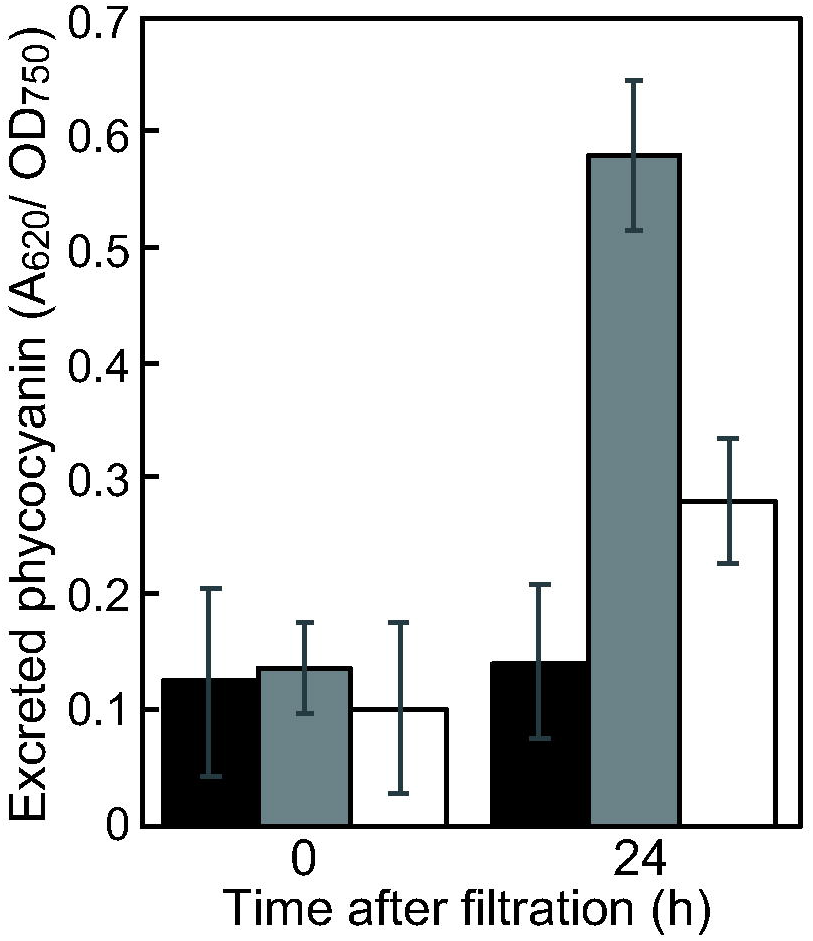
Excretion of phycocyanin after dehydration. Phycocyanin excreted in the medium from the cells of WT (black bars), the *orrA* disruptant (gray bars), and the *anaKa* disruptant (white bars) that were dehydrated on filters for 0 or 24 h was determined by measuring the absorbance at 620 nm.

## DISCUSSION

In this study, we revealed that the two-component response regulator OrrA played principal roles in the regulation of dehydration response and tolerance in *Anabaena. orrA* disruption abolished or diminished the induction of approximately 400 genes by dehydration (Fig.1 and Table S1). As previously indicated in response to salt stress (Ehira *et al*., 2014), OrrA regulated sucrose synthesis under dehydration conditions (Table 1). In addition, the trehalose synthesis genes *mth* and *mts*, which encode maltooligosyl trehalose hydrolase and maltooligosyl trehalose synthase, respectively, were regulated by OrrA (Table S1). Because disruption of *mth* and *mts* results in decreased dehydration tolerance (Higo *et al*., 2006), the depression of compatible solutes synthesis partly accounts for the susceptibility of the *orrA* disruptant to dehydration. In the OrrA regulon identified this study, *anaKa* was indispensable for dehydration tolerance in *Anabaena* (Fig. 5). *anaKa* encodes a protein of unknown function with a PRC-barrel domain at its N-terminus and a DUF2382 domain at its C-terminus. Although *anaKb, anaKc*, and *anaKd* were also induced by dehydration in an OrrA-dependent manner, their disruption did not affect dehydration tolerance. *anaKa* was more strongly induced than the other examined *anaK* genes (Fig. 2, Table S1). Differences in expression levels of AnaK proteins could explain the different phenotypes among the *anaK* mutants. OrrA and AnaK homologs are present in the genome of the terrestrial cyanobacteria, such as *Nostoc* sp. HK-01, *N. flagelliforme*, and *Chroococcidiopsis thermalis* PCC 7203. OrrA and AnaK homologs are highly expressed in desiccated samples in *N. flagelliforme* (Shang *et al*., 2019), and an AnaK homolog is induced by dehydration in *Nostoc* sp. HK-01 (Yoshimura *et al*., 2006), suggesting their conserved functions in desiccation tolerance.

The function of the PRC-barrel domain is unknown. It is found in the H subunit of photosystem reaction center (PRC) of the purple bacteria (Anantharaman and Aravind, 2002). The H subunit of PRC is necessary for functional PRC assembly, and it regulates electron transfer in PRC (Takahashi and Wraight, 1996; Lupo and Ghosh, 2004). The PRC-barrel domain is also found at the C-terminus of RimM, which is involved in RNA processing and ribosome assembly (Anantharaman and Aravind, 2002). The PRC-barrel domain of AnaKa is phylogenetically most closely related to that of the H subunit of PRC (Anantharaman and Aravind, 2002). AnaKa homologs are present in the highly radiation- and desiccation-resistant bacterium *Deinococcus radiodurans* and some species of the genus *Psychrobacter*, which are psychro- and osmo-tolerant bacteria (Kim *et al*., 2012). In *D. radiodurans*, the expression of the *anaKa* homolog DR1314 is upregulated by heat shock, and its induction is regulated by the extracytoplasmic function sigma factor Sig1 (Schmid *et al*., 2005). DR1314 is also regulated by the response regulator DrRRA, which is essential for radioresistance in *D. radiodurans* (Wang *et al*., 2016). The DR1314 disruptant is sensitive to hydrogen peroxide (H_2_O_2_) exposure and heat shock (Schmid *et al*., 2005; Wang *et al*., 2016). Thus, AnaK homologs play important roles in the stress resistance of extremophiles.

It was also indicated that OrrA regulates ROS defense systems (Table 2). ROS, which include singlet oxygen (^1^O_2_), superoxide anion (O_2_^−^), H_2_O_2_, and hydroxyl radical (•OH), is the main damaging factor during dehydration (Singh, 2018). O_2_^−^, which is generated by the reduction of oxygen through photosynthetic electron transport, is detoxified by superoxide dismutase (SOD), which dismutates it into H_2_O_2_ and O_2_ with. In *Anabaena, sodA* and *sodB* both encode SOD (Li *et al*., 2002), but their expression was not upregulated by dehydration. H_2_O_2_ is removed by catalases and peroxiredoxin (Prxs), which convert it into water. The catalase genes *katA* and *katB* were induced by dehydration; however, catalase activity is low in *Anabaena*, and Prxs play a more prominent role in H_2_O_2_ removal (Pascual *et al*., 2010). Although alr4641 (encoding 2-Cys-Prx), the overexpression of which protects *Anabaena* cells from oxidative stress (Banerjee *et al*., 2015), is induced by O_2^−^_ and H_2_O_2_ (Hurtado-Gallego *et al*., 2019), its expression was altered by dehydration. •OH is formed by the reaction of Fe^2+^ and H_2_O_2_, and Dps proteins sequestrate iron to prevent •OH formation (Howe *et al*., 2018). *Anabaena* carries four *dps* genes (all0458, all1173, alr3808, and all4145), and all0458 and all1173 were induced by dehydration. Thus, some H_2_O_2_-detoxifying enzymes were upregulated during dehydration, but H_2_O_2_ generation during dehydration is obscure because H_2_O_2_-responsive genes such as alr4641 and *trxA2* were not induced (Ehira and Ohmori, 2012; Hurtado-Gallego *et al*., 2019). Conversely, many genes encoding proteins involved in defense against ^1^O_2_ were induced by dehydration (Table 2). Excited chlorophylls can transfer excitation energy to oxygen, resulting in ^1^O_2_ formation (Latifi *et al*., 2009). High-light-inducible proteins, which are encoded by the *hli/scp* genes, prevent ^1^O_2_ formation by consuming the excess of PSII-generated electrons (Sinha *et al*., 2012). Orange carotenoid protein (OCP) is an efficient quencher of ^1^O_2_ (Sedoud *et al*., 2014). Recent findings indicated that All3221 and Alr4783 proteins of *Anabaena*, which are homologous to the N-terminal domain (NTD) of OCP, are effective quenchers of ^1^O_2_ (López-Igual *et al*., 2016). Moreover, in *N. flagelliforme*, a remarkable desiccation-tolerant terrestrial cyanobacterium, NTD-OCP proteins accumulated in cells collected from dried fields, suggesting its involvement in desiccation tolerance (Yang *et al*., 2019). AstaP, another OCP, was originally found in a green alga isolated from the dried surface of heated asphalt in midsummer, and it also displays ^1^O_2_ quenching activity (Kawasaki *et al*., 2013). *Anabaena* carries four genes encoding AstaP homologs (alr1819, all4647, all4894, and all5264), all of wchich were induced by dehydration. It is suggested that *N. commune* deactivates photosynthetic machinery in response to dehydration to protect photosystems from ^1^O_2_ generated through the imbalance between chlorophyll excitation and photosynthetic electron transport (Hirai *et al*., 2004). Thus, defense systems against ^1^O_2_ are especially important in dehydration tolerance of *Anabaena*.

**Table 2.**
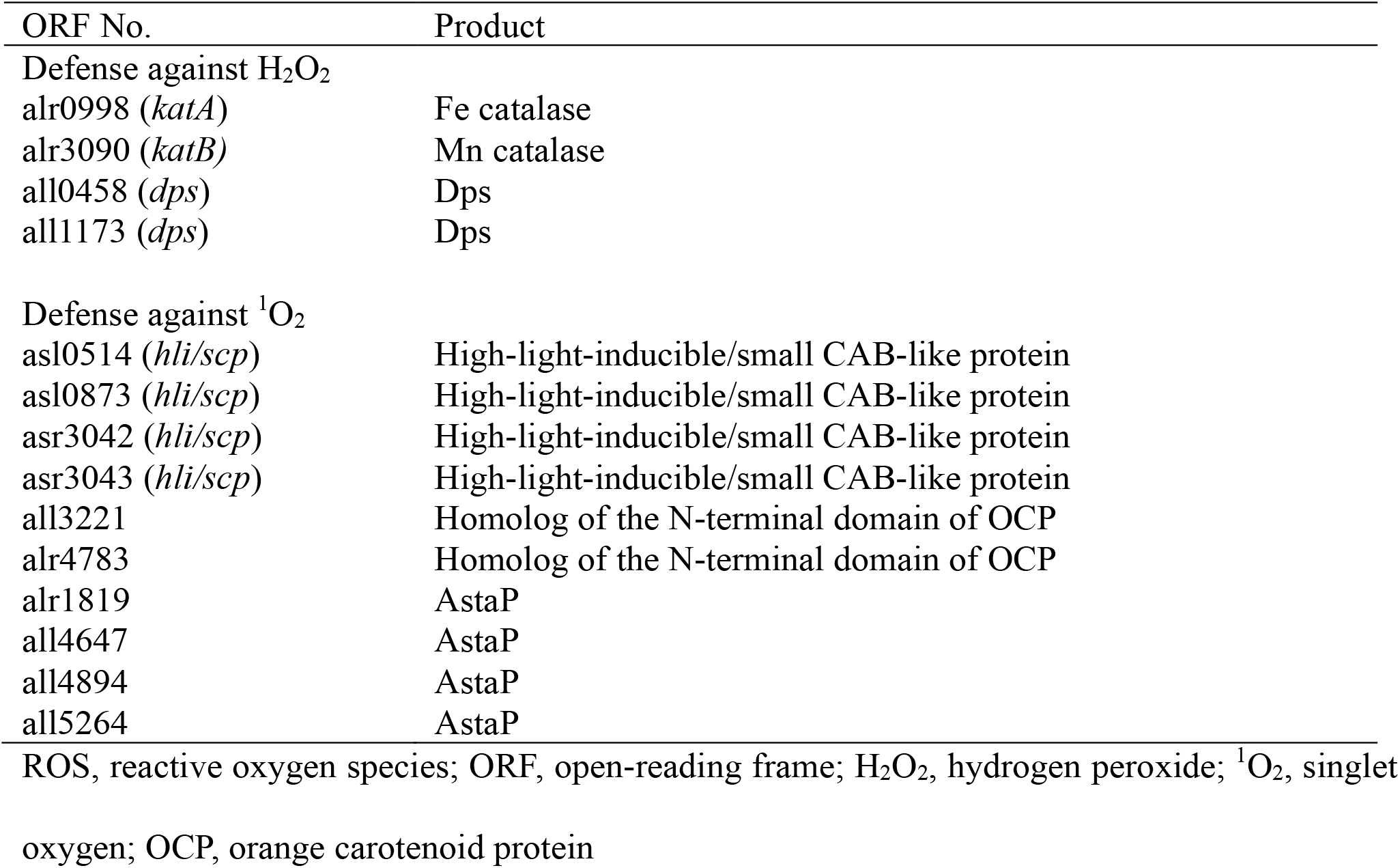
OrrA-regulated genes that are related to ROS defense

The OrrA regulon defined in this study contains a huge number of genes. A transcriptional network is likely to be involved in the dehydration response, because eight genes encoding transcriptional regulators, including *sigB2* (Ehira *et al*., 2014), were upregulated by dehydration under the control of OrrA (Table S1). In addition, the expression of more than 10 genes encoding signal transduction proteins, such as sensor histidine kinase (Hik) and Ser/The kinases, increased in an OrrA-dependent manner. These regulators constitute a regulatory network of the dehydration response of *Anabaena*.

OrrA was identified as a master regulator of the response to salt stress and dehydration and was involved in the response to low temperature (Figs. 1 and 4). OrrA was originally identified as the regulator of *lti2* in response to salt and osmotic response (Schwartz *et al*., 1998), implying that OrrA also plays an important role in the osmotic stress response. Evaporation of water from the medium surrounding cells increases external solute concentrations, resulting in salt and osmotic stresses. Therefore, a common signaling pathway could activate OrrA in response to dehydration, salt, and osmotic stresses. Because OrrA is a response regulator, its activity is regulated through the phosphorylation by a Hik. Three Hiks are located upstream of *orrA* in the genome of *Anabaena* (Ehira *et al*., 2014). These Hiks are involved in dehydration response, but none of them is a cognate Hik for OrrA in *Anabaena* (K. Shimonaka & S. Ehira, unpublished data). Identification of a cognate Hik for OrrA would reveal the mechanisms by which dehydration signals are sensed.

### EXPERIMENTAL PROCEDURES

#### Bacterial strains and culture conditions– Anabaena

sp. strain PCC 7120 and its derivatives were grown at 30°C under continuous illumination provided by a fluorescent lamp at 30 µE m^-2^ s^-1^ in the nitrogen-free modified Detmer’s medium (MDM_0_) (Watanabe, 1960). Liquid cultures were bubbled with air containing 1% (v/v) CO_2_. Salt stress conditions were generated by adding 50 mM NaCl to cultures in the late-logarithmic phase (optical density at 750 nm [OD_750_] = 0.5-0.7). Dehydration stress was imposed on *Anabaena* cells as described previously (Higo *et al*., 2006). A 25-ml portion of cultures in the late-logarithmic phase was filtered through 47-mm, 0.45-µm pore size mixed cellulose ester filters (Tokyo Roshi Kaisha, Tokyo, Japan) and dried for the indicated times at 30°C under continuous illumination provided by a fluorescent lamp at 10 µE m^-2^ s^-1^ in a Petri dish.

#### Mutant construction

All primers used in this study (Table S2) were designed based on genome data from CyanoBase (Fujisawa *et al*., 2017). DRorrAS was used as an *orrA* disruptant in this study (Ehira *et al*., 2014). DRanaKaK, DRanaKbK, DRanaKcS, and DRanaKdS, in which *anaKa* (*all4050*), *anaKb* (*all5315*), *anaKc* (*all4051*), and *anaKd* (*alr5332*), respectively, were disrupted, were constructed by replacing each gene with an antibiotic-resistant gene in the same manner as performed for DRorrAS using primer pairs specific for each gene (Ehira *et al*., 2014).

#### DNA microarray analysis

Total RNA was extracted from whole filaments according to Pinto et al. (Pinto *et al*., 2009) and was treated with DNase I (Takara Bio, Shiga, Japan). Global gene expression was analyzed using the *Anabaena* oligonucleotide microarray as described previously (Ehira and Ohmori, 2006). Microarray analyses were conducted using three sets of RNA samples isolated from independently grown cultures. Two hybridization reactions were performed with different combination of Cy-dyes for each set of RNA samples. Thus, six replicates were available per gene to determine changes in gene expression. Genes with at least twofold differences in transcript levels (*P* < 0.05; Student’s *t*-test) were identified. The microarray data were deposited in the KEGG Expression Database (accession numbers ex0001979 to ex0002002).

*qRT-PCR*– cDNAs were synthesized from 1 µg of total RNA with random hexamer primers using a PrimeScript first strand cDNA synthesis kit (Takara Bio). qRT-PCR was performed using the Thermal Cycler Dice Real Time System II (TP900; Takara Bio) in a 20-µl reaction mixture containing 10 µl of THUNDERBIRD SYBR qPCR Mix (Toyobo, Osaka, Japan), 0.2 µM each of gene-specific forward and reverse primers, and cDNA. Relative transcript levels were normalized to the value for 16S rRNA and represented as means of duplicate measurements. Experiments were repeated at least three times.

#### Dehydration tolerance assay

Cells dehydrated for the indicated times on filter paper were suspended in MDM_0_. The suspended cultures were diluted with MDM_0_ to OD_750_ of 0.1 and incubated at 30°C at 10 µE m^-2^ s^-1^. After 72 h, OD_750_ of the cultures was measured.

#### Determination of sucrose contents

The low-molecular-mass compounds of cells dehydrated for the indicated time were extracted with 80% ethanol for 3 h at 65°C (Higo *et al*., 2006). After centrifugation, supernatants were vacuum-dried, and the residue was dissolved in 0.5 ml of water. The samples were treated with 3 U of invertase (Sigma) in 50 mM sodium acetate (pH 4.5) for 2 h at 37°C, and the concentration of released glucose was determined using a Glucose Assay Kit (Biovision). The chlorophyll *a* content of cultures was determined as described previously (Mackinney, 1941).

#### Quantification of excreted phycobiliproteins

Dehydrated cells were resuspended in MDM_0_ and statically incubated for 5 min at room temperature. Cells were collected by centrifugation, and the absorbance of the supernatants at 620 nm was measured. A_620_ was normalized by OD_750_ measured prior to centrifugation.

## Acknowledgments

This work was supported by the Japan Society for the Promotion of Science (JSPS) [Challenging Research Exploratory 19K22290], and a grant from the Noda Institute of Scientific Research to SE.

## Author contributions

M. Ohmori and S. Ehira conceived the study. S. Kimura and S. Ehira designed the experiments. S. Kimura, M. Sato, X. Fan, and S. Ehira conducted the experiments and analyzed the data. S. Ehira wrote the manuscript, and all authors edited and approved the final manuscript.

## References

Adams, D.G., and Duggan, P.S. (1999) Heterocyst and akinete differentiation in cyanobacteria. New Phytol 144: 3–33.

Anantharaman, V., and Aravind, L. (2002) The PRC-barrel: a widespread, conserved domain shared by photosynthetic reaction center subunits and proteins of RNA metabolism. Genome Biol 3: 1–9.

Arii, S., Tsuji, K., Tomita, K., Hasegawa, M., Bober, B., and Harada, K.I. (2015) Cyanobacterial blue color formation during lysis under natural conditions. Appl Environ Microbiol 81: 2667–2675.

Banerjee, M., Chakravarty, D., and Ballal, A. (2015) Redox-dependent chaperone/peroxidase function of 2-Cys-Prx from the cyanobacterium Anabaena PCC7120: role in oxidative stress tolerance. BMC Plant Biol 15: 60.

Billi, D., Wright, D.J., Potts, M., Helm, R.F., Crowe, J.H., and Prickett, T. (2002) Engineering desiccation tolerance in Escherichia coli. Appl Environ Microbiol 66: 1680–1684.

Daly, M.J. (2009) A new perspective on radiation resistance based on Deinococcus radiodurans. Nat Rev Microbiol 7: 237–245.

Ehira, S., Kimura, S., Miyazaki, S., and Ohmori, M. (2014) Sucrose synthesis in the nitrogen-fixing cyanobacterium Anabaena sp. strain PCC 7120 is controlled by the two-component response regulator OrrA. Appl Environ Microbiol 80: 5672–5679.

Ehira, S., and Ohmori, M. (2006) NrrA, a nitrogen-responsive response regulator facilitates heterocyst development in the cyanobacterium Anabaena sp. strain PCC 7120. Mol Microbiol 59: 1692–1703.

Ehira, S., and Ohmori, M. (2012) The redox-sensing transcriptional regulator RexT controls expression of thioredoxin A2 in the cyanobacterium Anabaena sp. strain PCC 7120. J Biol Chem 287: 40433–40440.

Ehira, S., Ohmori, M., and Sato, N. (2005) Identification of low-temperature-regulated ORFs in the cyanobacterium Anabaena sp. strain PCC 7120: distinguishing the effects of low temperature from the effects of photosystem II excitation pressure. Plant Cell Physiol 46: 1237–1245.

Fagliarone, C., Mosca, C., Ubaldi, I., Verseux, C., Baqué, M., Wilmotte, A., and Billi, D. (2017) Avoidance of protein oxidation correlates with the desiccation and radiation resistance of hot and cold desert strains of the cyanobacterium Chroococcidiopsis. Extremophiles 21: 981–991.

Flores, E., Picossi, S., Valladares, A., and Herrero, A. (2019) Transcriptional regulation of development in heterocyst-forming cyanobacteria. Biochim Biophys Acta Gene Regul Mech 1862: 673–684.

França, M.B., Panek, A.D., and Eleutherio, E.C. (2007) Oxidative stress and its effects during dehydration. Comp Biochem Physiol A Mol Integr Physiol 146: 621–631.

Fredrickson, J.K., Li, S.M., Gaidamakova, E.K., Matrosova, V.Y., Zhai, M., Sulloway, H.M., et al. (2008) Protein oxidation: key to bacterial desiccation resistance? ISME J 2: 393–403.

Fujisawa, T., Narikawa, R., Maeda, S.I., Watanabe, S., Kanesaki, Y.Y., Kobayashi, K., et al. (2017) CyanoBase: a large-scale update on its 20th anniversary. Nucleic Acids Res 45: D551–D554.

Hershkovitz, N., Oren, A., and Cohen, Y. (1991) Accumulation of trehalose and sucrose in cyanobacteria exposed to matric water stress. Appl Environ Microbiol 57: 645–648.

Higo, A., Katoh, H., Ohmori, K., Ikeuchi, M., and Ohmori, M. (2006) The role of a gene cluster for trehalose metabolism in dehydration tolerance of the filamentous cyanobacterium Anabaena sp. PCC 7120. Microbiology 152: 979–987.

Hirai, M., Yamakawa, R., Nishio, J., Yamaji, T., Kashino, Y., Koike, H., and Satoh, K. (2004) Deactivation of photosynthetic activities is triggered by loss of a small amount of water in a desiccation-tolerant cyanobacterium, Nostoc commune. Plant Cell Physiol 45: 872–878.

Howe, C., Ho, F., Nenninger, A., Raleiras, P., and Stensjö, K. (2018) Differential biochemical properties of three canonical Dps proteins from the cyanobacterium Nostoc punctiforme suggest distinct cellular functions. J Biol Chem 293: 16635–16646.

Hurtado-Gallego, J., Redondo-López, A., Leganés, F., Rosal, R., and Fernández-Piñas, F. (2019) Peroxiredoxin (2-cys-prx) and catalase (katA) cyanobacterial-based bioluminescent bioreporters to detect oxidative stress in the aquatic environment. Chemosphere 236: 124395.

Katoh, H., Shiga, Y., Nakahira, Y., and Ohmori, M. (2003) Isolation and characterization of a drought-tolerant cyanobacterium, Nostoc sp. HK-01. Microbes Environ 18: 82–88.

Kawasaki, S., Mizuguchi, K., Sato, M., Kono, T., and Shimizu, H. (2013) A novel astaxanthin-binding photooxidative stress-inducible aqueous carotenoprotein from a eukaryotic microalga isolated from asphalt in midsummer. Plant Cell Physiol 54: 1027–1040.

Kim, S.J., Shin, S.C., Hong, S.G., Lee, Y.M., Choi, I.G., and Park, H. (2012) Genome sequence of a novel member of the genus Psychrobacter isolated from Antarctic soil. J Bacteriol 194: 2403–2403.

Latifi, A., Ruiz, M., and Zhang, C.C. (2009) Oxidative stress in cyanobacteria. FEMS Microbiol Rev 33: 258–278.

Li, T., Huang, X., Zhou, R., Liu, Y., Li, B., Nomura, C., and Zhao, J. (2002) Differential expression and localization of Mn and Fe superoxide dismutases in the heterocystous cyanobacterium Anabaena sp. strain PCC 7120. J Bacteriol 184: 5096–5103.

Lipman, C.B. (1941) The successful revival of Nostoc commune from a herbarium specimen eighty-seven years old. Bull Torrey Bot Club 68: 664.

López-Igual, R., Wilson, A., Leverenz, R.L., Melnicki, M.R., Bourcier de Carbon, C., Sutter, M., et al. (2016) Different functions of the paralogs to the N-terminal domain of the orange carotenoid protein in the cyanobacterium Anabaena sp. PCC 7120. Plant Physiol 171: 1852–1866.

Lupo, D., and Ghosh, R. (2004) The reaction center H subunit is not required for high levels of light-harvesting complex 1 in Rhodospirillum rubrum mutants. J Bacteriol 186: 5585–5595.

Mackinney, G. (1941) Absorption of light by chlorophyll solutions. J Biol Chem 140: 315–322.

Ohmori, M., Ikeuchi, M., Sato, N., Wolk, P., Kaneko, T., Ogawa, T., et al. (2001) Characterization of genes encoding multi-domain proteins in the genome of the filamentous nitrogen-fixing cyanobacterium Anabaena sp. strain PCC 7120. DNA Res 8: 271–284.

Pascual, M.B., Mata-Cabana, A., Florencio, F.J., Lindahl, M., and Cejudo, F.J. (2010) Overoxidation of 2-Cys peroxiredoxin in prokaryotes: cyanobacterial 2-Cys peroxiredoxins sensitive to oxidative stress. J Biol Chem 285: 34485–34492.

Pinto, F.L., Thapper, A., Sontheim, W., and Lindblad, P. (2009) Analysis of current and alternative phenol based RNA extraction methodologies for cyanobacteria. BMC Mol Biol 10: 79.

Pointing, S.B., and Belnap, J. (2012) Microbial colonization and controls in dryland systems. Nat Rev Microbiol 10: 551–562.

Potts, M. (2001) Desiccation tolerance: a simple process? Trends Microbiol 9: 553–559.

Sakamoto, T., Yoshida, T., Arima, H., Hatanaka, Y., Takani, Y., and Tamaru, Y. (2009) Accumulation of trehalose in response to desiccation and salt stress in the terrestrial cyanobacterium Nostoc commune. Phycol Res 57: 66–73.

Schmid, A.K., Howell, H.A., Battista, J.R., Peterson, S.N., and Lidstrom, M.E. (2005) Global transcriptional and proteomic analysis of the Sig1 heat shock regulon of Deinococcus radiodurans. J Bacteriol 187: 3339–3351.

Schwartz, S.H., Black, T.A., Jäger, K., Panoff, J.M., and Wolk, C.P. (1998) Regulation of an osmoticum-responsive gene in Anabaena sp. strain PCC 7120. J Bacteriol 180: 6332–6337.

Sedoud, A., López-Igual, R., Ur Rehman, A., Wilson, A., Perreau, F., Boulay, C., et al. (2014) The cyanobacterial photoactive orange carotenoid protein is an excellent singlet oxygen quencher. Plant Cell 26: 1781–1791.

Shang, J.L., Chen, M., Hou, S., Li, T., Yang, Y.W., Li, Q., et al. (2019) Genomic and transcriptomic insights into the survival of the subaerial cyanobacterium Nostoc flagelliforme in arid and exposed habitats. Environ Microbiol 21: 845–863.

Singh, H. (2018) Desiccation and radiation stress tolerance in cyanobacteria. J Basic Microbiol 58: 813–826.

Singh, H., Anurag, K., and Apte, S.K. (2013) High radiation and desiccation tolerance of nitrogen-fixing cultures of the cyanobacterium Anabaena sp. strain PCC 7120 emanates from genome/proteome repair capabilities. Photosynth Res.

Sinha, R.K., Komenda, J., Knoppová, J., Sedlářová, M., and Pospíšil, P. (2012) Small CAB-like proteins prevent formation of singlet oxygen in the damaged photosystem II complex of the cyanobacterium Synechocystis sp. PCC 6803. Plant Cell Environ 35: 806–818.

Takahashi, E., and Wraight, C.A. (1996) Potentiation of proton transfer function by electrostatic interactions in photosynthetic reaction centers from Rhodobacter sphaeroides: first results from site-directed mutation of the H subunit. Proc Natl Acad Sci U S A 93: 2640–2645.

Tamaru, Y., Takani, Y., Yoshida, T., and Sakamoto, T. (2005) Crucial role of extracellular polysaccharides in desiccation and freezing tolerance in the terrestrial cyanobacterium Nostoc commune. Appl Environ Microbiol 71: 7327–7333.

Tapia, H., and Koshland, D.E. (2014) Trehalose is a versatile and long-lived chaperone for desiccation tolerance. Curr Biol 24: 2758–2766.

Tibiletti, T., Rehman, A.U., Vass, I., and Funk, C. (2018) The stress-induced SCP/HLIP family of small light-harvesting-like proteins (ScpABCDE) protects photosystem II from photoinhibitory damages in the cyanobacterium Synechocystis sp. PCC 6803. Photosynth Res 135: 103–114.

Wang, L., Hu, J., Liu, M., Yang, S., Zhao, Y., Cheng, K., et al. (2016) Proteomic insights into the functional basis for the response regulator DrRRA of Deinococcus radiodurans. Int J Radiat Biol 92: 273–280.

Wang, L., Sun, Y.P., Chen, W.L., Li, J.H., and Zhang, C.C. (2002) Genomic analysis of protein kinases, protein phosphatases and two-component regulatory systems of the cyanobacterium Anabaena sp. strain PCC 7120. FEMS Microbiol Lett 217: 155–165.

Watanabe, A. (1960) List of algal strains in collection at the Institute of Applied Microbiology, University of Tokyo. J Gen Appl Microbiol 6: 283–292.

Xu, H.F., Dai, G.Z., Ye, D.M., Shang, J.L., Song, W.Y., Shi, H., and Qiu, B.S. (2020) Dehydration-induced DnaK2 chaperone is involved in PSII repair of a desiccation-tolerant cyanobacterium. Plant Physiol 182: 1991–2005.

Yang, Y.W., Yin, Y.C., Li, Z.K., Huang, D., Shang, J.L., Chen, M., and Qiu, B.S. (2019) Orange and red carotenoid proteins are involved in the adaptation of the terrestrial cyanobacterium Nostoc flagelliforme to desiccation. Photosynth Res 140: 103–113.

Yoshimura, H., Ikeuchi, M., and Ohmori, M. (2006) Up-regulated gene expression during dehydration in a terrestrial cyanobacterium, Nostoc sp. strain HK-01. Microbes Environ 21: 129–133.

Yoshimura, H., Okamoto, S., Tsumuraya, Y., and Ohmori, M. (2007) Group 3 sigma factor gene, sigJ, a key regulator of desiccation tolerance, regulates the synthesis of extracellular polysaccharide in cyanobacterium Anabaena sp. strain PCC 7120. DNA Res 14: 13–24.

Zhou, R., and Wolk, C.P. (2002) Identification of an akinete marker gene in Anabaena variabilis. J Bacteriol 184: 2529–2532.

