## Supplementary material for "The two-component response regulator OrrA confers dehydration tolerance by regulating *anaKa* expression in the cyanobacterium *Anabaena* sp. strain PCC 7120": Table S2

Table S2. Primers used in this study

| Primer | Sequence (5′–3′) |
|---|---|
| Primers for mutant construction | |
| anaKa-5F | AAGAGCTCAAAGTGGCGTGAGAGTTGG |
| anaKa-5R | AAGGATCCTGTAAAGTGCCATGTGATTCCA |
| anaKa-3F | AAGGATCCACGTGAATGCTCCTGGTCTT |
| anaKa-3R | AACTCGAGTGGTCACGCTAGCCATAAA |
| anaKb-5F | AAGAGCTCGCAGCCATTTATTCAGGGT |
| anaKb-5R | AAGGATCCAGTCGGCAGCAAACTCTTCT |
| anaKb-3F | AAGGATCCACAGTTGAAGCCGAAGAGAC |
| anaKb-3R | AACTCGAGAATCTCCTGGGCTGGGTAAC |
| anaKc-5F | AAGAGCTCCGAAGCTTACCCAGAAAGA |
| anaKc-5R | AAGGATCCTCTGGCTGACGACTGAAGT |
| anaKc-3F | AAGGATCCACTTGCTTGCTTTCTGGAGT |
| anaKc-3R | AACTCGAGCGTTCTCTTTCCACGGGTAC |
| anaKd-5F | AAGAGCTCGGTGGATTCAAGGGTCATCC |
| anaKd-5R | AAGGATCCTCATTGTCAGCGTAAACATCAA |
| anaKd-3F | AAGGATCCTTACTTCAGAACATACGCA |
| anaKd-3R | AACTCGAGGGAATTGTAATCACCGCAAG |
| Primers for qRT-PCR | |
| RTrrn16S-F2 | GCAAGTCGAACGGTCTCTTC |
| RTrrn16S-R2 | GGTATTAGCCACCGTTTCCA |
| RTorrA-F | GGGCTAAGAGCTGCGTTACA |
| RTorrA-R | CGGTATGATCCATTGTCAGG |
| RTmth-F | AATTGGCGCTCACTACTTGG |
| RTmth-R | TTCCTCTGGCTTTAAGATTTGC |
| RT0458-F | CCTCAAAGAACACGCTGGAG |
| RT0458-R | CTGCGGTCTACATGGTCTCT |
| RT1173-F | GTCTCTATCCCACCCGCATT |
| RT1173-R | GAGTGGCCGCTAAAGTTTGA |
| RTanaKa-F | TACCCGAATTCAACGACCGT |
| RTanaKa-R | GTTGATCTGTCTGCGCTTGT |
| RTanaKb-F | AAGGTGCGTGGTGTTTATCG |
| RTanaKb-R | CGATCGTGAGCAACTGAAGG |
| RTanaKc-F | TGTTGACGGACTGAGCAAAG |
| RTanaKc-R | CACCGCGTACTCTTTCTTCG |
| RTanaKd-F | TGTTGGTACCTGGGCTAACT |
| RTanaKd-R | CCTGGCGTTGCTAATGGTAC |
